# Pseudo batch transformation: A novel method to correct for mass removal through sample withdrawal of fed-batch fermentations

**DOI:** 10.1101/2024.05.27.596043

**Authors:** Viktor Hesselberg-Thomsen, Teddy Groves, Tim McCubbin, Iván Martínez-Monge, Igor Marín de Mas, Lars Keld Nielsen

## Abstract

**Summary:** We present a novel “pseudo batch” transformation algorithm that maps analytical data obtained for fed-batch bioreactor cultivations onto a constant volume batch process, significantly decreasing the complexity of characterizing the fed-batch process.

**Availability and implementation:** Our method is implemented in both Excel and Python and is available with tutorials and example data from https://github.com/biosustain/pseudobatch. The Python package is also available on PYPI under the name “pseudobatch”.

**Contact:** Lars Keld Nielsen

**Supplementary information:** A comprehensive explanation of the simulated fed-batch, parameter estimation procedures, and the Bayesian model can be found in supplementary information (S1-S5).

## Introduction

Fed-batch cultivation is commonly used by the biomanufacturing industry to avoid substrate inhibition and/or overflow metabolism in both mammalian and microbial systems [1]. The analysis of fed-batch data is confounded by the dynamic changes in bioreactor volume caused by sampling and feeding processes. The removal of mass through sampling leads to discontinuous mass time-evolution curves (Figure 1A), which will lead to inaccurate rate, yield, and metabolic flux estimates and inaccurate mass balancing unless carefully considered. Furthermore, this limits the ability of machine learning algorithms to train on fed-batch data.

**Figure 1.**
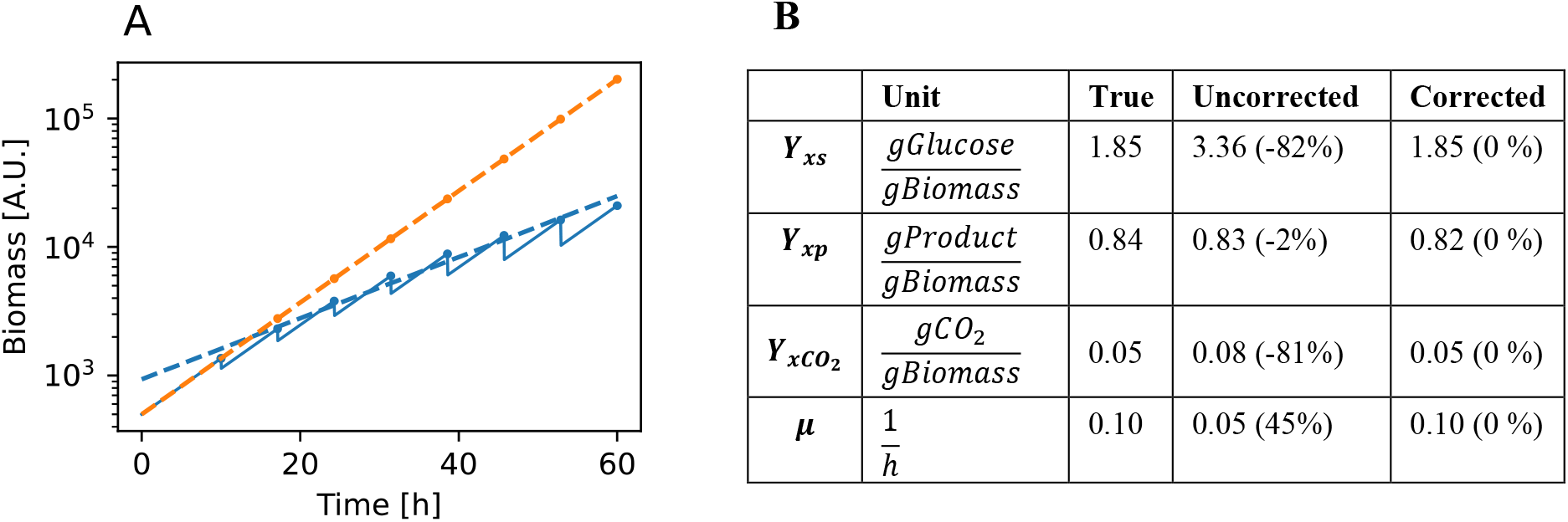
**A)** Simulated exponential fed-batch with sampling (blue line), sample measurements (blue dots) and the pseudo batch transform measurements (orange dots). Fitted growth rate model for raw measurements (blue dashed line) and pseudo batch transformed measurements (orange dashed line). **B)** Uncorrected and corrected estimated values of the yields and growth rate (see S4 for estimation approach). The parenthesis holds the percent error of the estimate compared to the true value.

Traditionally, fed batch processes have been optimized in larger bioreactors where the sample fraction and hence the effect is negligible. However, high-throughput screening for clone selection or feeding strategies is increasingly performed using miniaturized multiplex fermentation systems (e.g., ambr15, BioLector) [2]. Rapid advances in the use of machine learning for optimizing bioprocesses is one of the drivers for high-throughput small volume cultivation [3], [4] and most systems suffer from substantial sampling fraction. Even reactors with larger operating volumes can be subject to substantial errors when significant samples are withdrawn to prevent reactor overflow, extend cultivation time and reduce time between fermentations [5], [6], particularly in the case where reactors are operated as cyclic or repetitive fed-batch modes.

In this study, we present a novel method called pseudo batch transformation that removes the effects of the volume changes from the experimental data. We apply this method to simulated data to illustrate that it estimates the correct rates and yields using standard linear modeling procedures [7].

## Methods

### Formalization of the Pseudo batch transformation method

The pseudo batch transformation independently projects time-course data for each species (biomass or metabolite) measured across a fed-batch bioreactor culture onto a constant volume batch culture. For convenience, we define an accumulated dilution factor (ADF), defined in equation (1), which scales a concentration at any given measurement time to the original bioreactor volume.

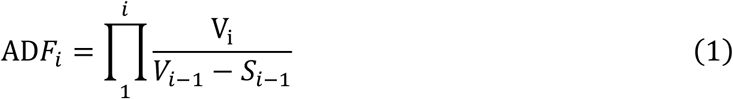

We employ an integer-valued time index variable *i* which denotes the order of the measurements. S_i−1_ is the sample volume at time index *i* − 1. This is 0 at time points where no sample was withdrawn. *V*_*i*_ is the volume prior to sampling at time index *i*. We see that *ADF*_*i*_ = 1 when no volume is added to the reactor, i.e. no dilution.

The pseudo batch transformation is defined by equation (2):

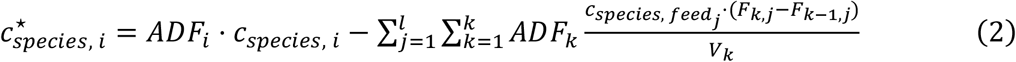

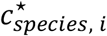 is the pseudo batch concentration at measurement time index *i*, which depends on two terms. The first term scales the measured concentration of a species at time index *i, c*_*species, i*_, by the accumulated dilution factor at time index *i, ADF*_*i*_ . This term essentially transforms a measured concentration to the concentration that would be achieved if the bioreactor was not being diluted by the feed. The second term accumulates species added through feeding. Essentially, this term performs a scaled integration from the beginning until the *i*’th measurement by iterating *k* from 1 until *i*. The concentration of the added species at each time index, *k*, is scaled by its corresponding ADF at time index *k*, across all *l* feeds. 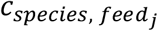 is the concentration of the species in the feed *j, F*_*k*,*j*_ is the accumulated volume of feed *j* added at time *k* and *V*_*k*_ is the volume at time index *k* prior to sampling. In the case where the species is not fed, i.e. a product, this second term collapses to 0.

The pseudo batch concentrations represent the effective concentration of catabolized substrates or generated products and can be directly utilized within yield calculations. The method can be expanded to capture substrates or products that are not solubilized in the media, such as gas data, by converting these species into a pseudo concentration (see S2 for details). Furthermore, pseudo concentrations are allowed to be negative. Importantly, they should not be interpreted as an actual prediction of the theoretical fed-batch process without volume changes. This would likely result in a different environment in the bioreactor, e.g., due to larger cell densities. Rather the method scales the observed phenotypic behavior to be more interpretable both visually and through statistical models.

## Results and discussion

### Assessing the validity of the pseudo batch method with simulated data

To illustrate the need and utility of pseudo batch transformation, we simulated a microbial culture under fed-batch conditions. In this simulation glucose is consumed, and biomass, product, and *CO*_2_ are produced. The feed addition follows an exponential feeding profile designed to keep the substrate concentration constant (further detailed in S3). When samples are withdrawn from this culture, mass is removed, and the true mass time course becomes discontinuous (Figure 1A - blue line and figure S1). Thus, if we estimate the growth rate as the slope of a log-linear model, we obtain a biased estimate which is 34% lower than the simulated growth rate (Figure 1A – dashed blue line and B). However, if the pseudo batch transformation is applied to the data before estimating the growth rate, the correct values are obtained (Figure 1A – dashed orange line). The same issue applies to the yield calculations without the pseudo batch transformation (Figure 1B), except where a yield is calculated between two products which cancels out the majority of the error. Often the production or consumption rates are of interest, the calculation of which will compound the error from the yields and growth rate estimates.

### The pseudo batch transformation introduces negligible additional error

The pseudo batch concentrations are calculated from several measured quantities: concentration, bioreactor volume, feeding volume, and concentrations in the feed medium. Uncertainty in these quantities will combine to produce uncertainty in the pseudo batch concentration. We therefore created a Bayesian statistical model that propagates known uncertainties and sources of measurement error into uncertainties for pseudo batch transformed concentrations. We fit this model for a typical case under a plausible range of error assumptions, assuming independent 5% errors on reactor volume, sample volume measurements and a consistent roughly +/-10% error in feed volume delivered.

In our case study, the logarithmic-scale uncertainty, as captured by the marginal coefficients of variation of posterior samples, was approximately the same for transformed and untransformed concentrations under a range of plausible measurement error assumptions (Figure S4). This indicates that, at least in such a typical case, the pseudo batch transformation introduces negligible additional error. We also used these posterior samples to recover known growth rates and yield coefficients using standard methods for batch fermentation data. This demonstrates, again for this typical case, that analyses using this data are appropriately accurate (Figure S5 and Table S3).

Further details on the model and results can be found in Section S5. Additionally, we exposed our model in the package through a function. This allow users to quantify their own pseudo batch transformation uncertainties (Further detailed in https://biosustain.github.io/pseudobatch/Tutorials/7%20-%20Pseudobatch%20transformation%20with%20uncertainties.html).

### The pseudo batch transformation enables the utilization of an extensive toolkit

The pseudo batch transformed data can be used in the standard linear model workflows in Excel, R, Python or other software, to estimate rates and yields from experimental measurements. Furthermore, the continuous nature of the pseudo batch transformed data enables a big suite of more advanced statistical and machine learning methods to work straight out of the box, which eases data integration. For example the pseudo batch transformation enables estimation of rates through gaussian process regression [8], spline fitting [9], and dynamic flux balance analysis [9]. Additionally, our method enables training of machine learning algorithms on data from sampled microbioreactor cultivations. Such algorithms could include physics-informed neural networks for bioprocess modelling [10] or strain optimization through a probabilistic ensemble model [4]. Machine learning methods are crucial techniques for high throughput data processing pipelines and automated decision making [3]. The pseudo batch transformation enables seamless use of fed-batch data from microbioreactor systems in such pipelines.

## Conclusion

The pseudo batch transformation method presented in this study effectively transforms measurements from sampled fed-batch cultivations into a space where it is possible to calculate rates and yields using standard methods. The transformation does not increase the uncertainty of the measurements or estimated values. The pseudo batch transformation can facilitate data-driven decision making and strain characterization based on fed-batch fermentations, which can enhance bioprocess optimization, clone selection, and metabolic studies and enables integration of fed-batch data with machine learning models in the biotechnology industry and academia.

## Acknowledgements

None

