## Supplementary information for "Pseudo batch transformation: A novel method to correct for mass removal through sample withdrawal of fed-batch fermentations"

#### S1 Code and reproduction

All Python required to reproduce the results can be found in our Github repository (*https://github.com/biosustain/pseudobatch* (*DOI:10.11583/DTU.23453018*)). This also contains a Docker image which can be used to reproduce the environment used to run simulations and analysis.

#### S2 Pseudo batch transformation of gaseous species

Gaseous species are either not or only partially affected by sample withdrawal, therefore they require special treatment when using the pseudo batch transformation. To derive the special treatment for gaseous species we start with at a general mass balance over a bioreactor for any one species.

$$\frac{dM_{species}(t)}{dt} = In_{gas,\,species}(t) + In_{liquid,\,species}(t) - Out_{gas,\,species}(t) + Metabolism_{species}(t) - Sampled_{species}(t)$$

where $M_{species}$ is the total amount of species in the reactor in both gas and aqueous phase. The term $Metabolism_{species}$ is the net result of the microbial metabolism and covers both production (positive) and consumption (negative). The $In$ variables describe the amounts of species entering the bioreactor through inlet stream of gas or liquid. The $Out_{gas,\,species}$ describes the amounts of species leaving the bioreactor through outlet gas stream. We only consider fed-batch operations, thus a liquid outlet stream is omitted. Finally, the term $Sampled_{species}$ describes the amounts of species removed through sampling. All the terms are functions over time, $t$. If we integrate the mass balance, we get the following:

$$M_{species}(t) = \int_0^t In_{gas,\,species}(t)dt + \int_0^t In_{liquid,\,species}(t)dt - \int_0^t Out_{gas,\,species}(t)dt + \int_0^t Metabolism_{species}(t)dt - \int_0^t Sampled_{species}(t)dt$$

This shows that the concentration of the species at a given time is a result of the integrated mass balance, i.e. the accumulated "events", e.g. input of medium, gas, sampling etc. We are interested in the contribution of the microbes to the mass balance thus we will isolate the metabolism term.

$$\int_0^t Metabolism_{species}(t)dt = M_{species}(t) - \int_0^t In_{gas,\,species}(t)dt + \int_0^t In_{liquid,\,species}(t)dt + \int_0^t Out_{gas,\,species}(t)dt + \int_0^t Sampled_{species}(t)dt$$

In a real-world setting, we do not have access to the values of these integrals. Instead, we have measurements which estimate the terms. The liquid concentration, $M_{species}(t)$ is commonly measured through light spectrometry, HPLC or mass spectrometry methods. Gaseous species can occasionally be measured using probes or is otherwise estimated using gas-liquid equilibrium relationships such as Henry's law. The inlet integrals are estimated through flow rates and known concentrations in inlet gas and liquid. The outlet integral is estimated using an off-gas measuring device and the mass removal due to sampling can be calculated from the sample volume and the liquid concentration measurement. All these are discrete measurements which estimates each integral at a specific time point. For the rest of this section, we will use measured integral values which we will denote as **Accumulated quantities**. To indicate that these contain discrete values we will use t in square brackets, $[t]$, to denote the value at a specific time point. In this discrete notation the mass balance looks as follows.

$$AccumMetabolism_{species}[t] = M_{species}[t] - AccumIn_{gas,\,species}[t] + AccumIn_{liquid,\,species}[t] + AccumOut_{gas,\,species}[t] + AccumSampled_{species}[t]$$

The pseudo batch Python package provide a utility function which calculate the accumulated metabolism using the mass balance above. The function is called $metabolised\_amount$.

#### S2.1 Hypothetical concentration

Before applying the pseudo batch transformation to the measurements of the gaseous species, the measurements need some preprocessing. To preprocess the fermentation data, we calculate the *hypothetical concentration* of the species if it were fully solubilised in the media inside the bioreactor. Hypothetical concentration takes into account that *if species in a gaseous form was more solubilised, then more of the species would be removed through sample withdrawal*. We calculate the hypothetical concentration using the following equation.

$$Hypothetical\ concentration_{species}[t] = (AccumMetabolism[t] - AccumSampleLoss[t]) * (V[t] - Sample\ volume[t])^{-1}$$

Where

$$AccumSampleLoss[t] = AccumSampleLoss[t-1] + \sum_{1}^{t}(AccumMetabolism[t] - AccumSampleLoss[t-1]) * \frac{Sample\ volume[t]}{V[t]}$$

The calculation of the hypothetical concentration is provided by the Python function $hypothetical\_concentration$. The hypothetical concentrations can be transformed through the pseudo batch transformation. To see examples of this please see the *Special case: gaseous species tutorial* in our documentation.

### S3 Simulated data

To validate the pseudo batch transformation method, we simulated bioreactors in different scenarios standard exponential fed-batch, exponential fed-batch with product inhibition, exponential fed-batch with volatile product, and fed-batch with multiple discrete feeds. All simulations were set up as systems of ordinary differential equations in Julia. We used the DifferentialEquation.jl package to solve the equation systems [1]. In the paper we only present data and estimated quantities from the standard exponential fed-batch simulation. In the following we provide a description of standard exponential fed-batch simulation. To see details of the other simulations we refer the reader to the code in the Github repository (*https://github.com/biosustain/pseudobatch/tree/main/article/simulation_scripts*).

The standard exponential fed-batch simulation considers a microbial culture which consumes glucose and produces biomass, a generic product and $CO_2$ according to the following system of ordinary differential equations:

$$\frac{dm_s}{dt} = -Yxs * \mu(t) * m_x + v_{feed}(t, F0, \mu_0) * c_f$$

$$\frac{dm_x}{dt} = \mu(t) * m_x$$

$$\frac{dm_p}{dt} = Yxp * \mu(t) * m_x$$

$$\frac{dm_{co2}}{dt} = Yxco2 * \mu(t) * m_x$$

$$\frac{dV}{dt} = v_{feed}(t, F_0, \mu_0)$$

Where $Y_{xs}$, $Y_{xp}$, and $Y_{xco2}$ are the biomass yield coefficients, in $\frac{g}{g}$, of glucose, a generic product and $CO_2$; $m_s$, $m_p$, and $m_{co2}$ are the mass in grams of glucose, a generic product, and $CO_2$, respectively; $c_f$ is the concentration of glucose in the feed medium and $v_{feed}(t)$ is the feeding profile, i.e., the flow rate of the feed medium at time $t$.

The growth rate was modelled through Monod kinetics [2]:

$$\mu(t) = \mu_{max} \frac{c_s(t)}{c_s(t) + K_s}$$

The feeding profile is calculated to achieve a constant specific growth rate through the following equation [2]:

$$v_{feed}(t, F_0, \mu_0) = F_0 * e^{\mu_0 t}$$

To simplify the testing and validation, we used the exact values for $Y_{sx}$ and $X_0$ when calculating the feeding profile, even though these would not be exactly known in a true experimental setup.

We simulated samples of the fed-batch process using the callback functionality in DifferentialEquations.jl. These were activated at specific time points. A given volume was removed from the simulated bioreactor, and with that also mass of the species in the culture. The feeding profile was then adjusted to accommodate the decrease in total biomass. This adjustment is required to maintain a constant specific growth rate. The adjusted feeding profile was calculated as follows:

$$F_0^{adjusted} = F_0^{original} * \frac{V_{after\ sampling}}{V_{before\ sampling}}$$

$$v_{feed}^{adjusted}(t) = F_0^{adjusted} * e^{\mu_0 t}$$

The growth parameters were set to textbook estimates for *Saccharomyces cerevisiae* from [2]. To avoid an initial adaptation phase in the growth rate, we calculated the glucose concentration resulting in the desired growth rate by isolating $c_s(t)$ Monod kinetics equation.

The standard exponential fed-batch simulation is plotted in figure S1 and the concentrations are visualized along with pseudo batch transformed concentrations in figure S2.

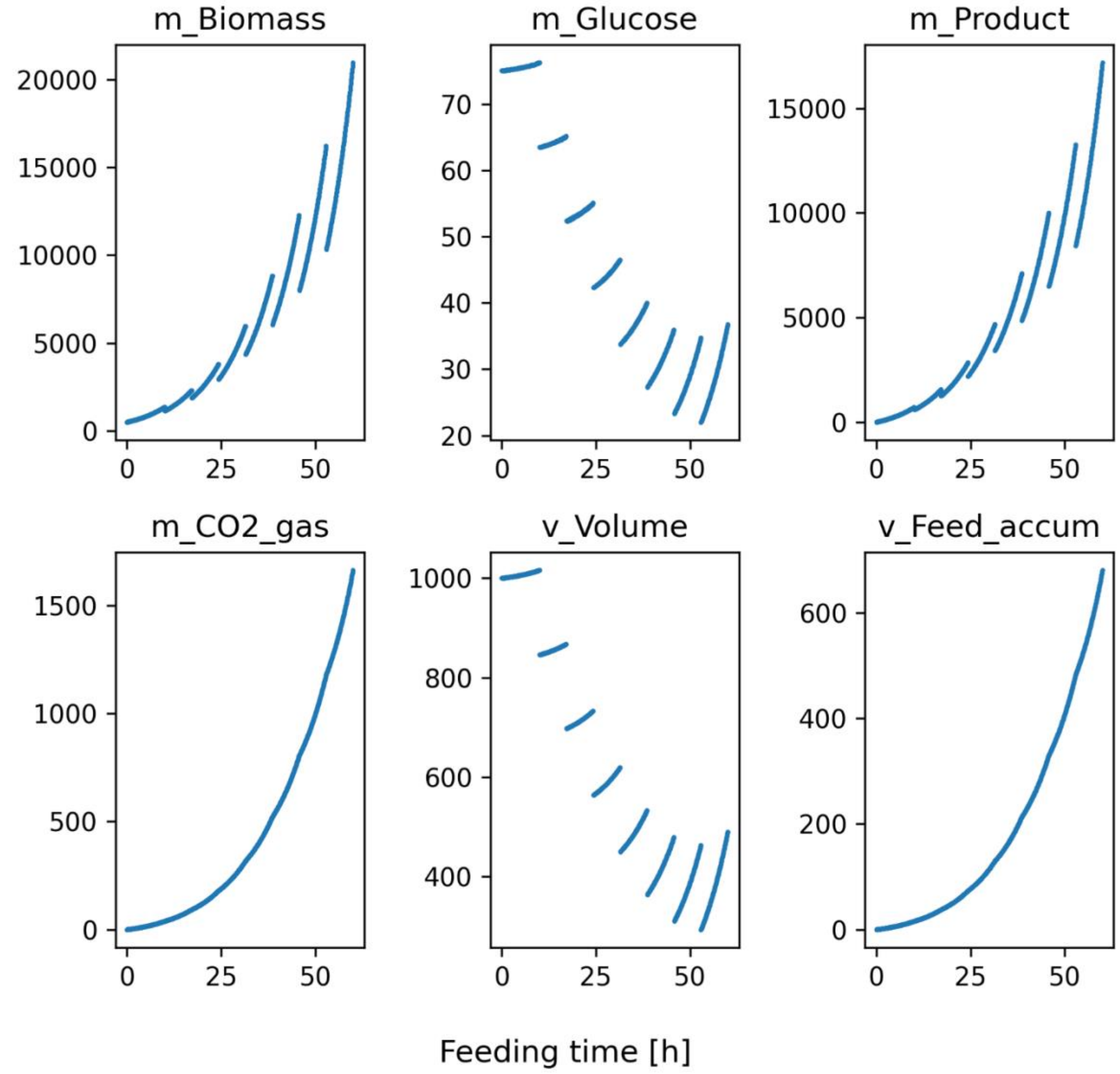

***Figure S1*** *standard exponential fed-batch simulated data. Panels m_Biomas, m_Glucose, m_Product show the total mass time series for biomass, glucose, and product, respectively. m_CO2_gas shows the mass of CO2 leaving the bioreactor, mimicking an off-gas measurement. v_Volume is the bioreactor volume and v_Feed_accum is the accumulated volume of feed added.*

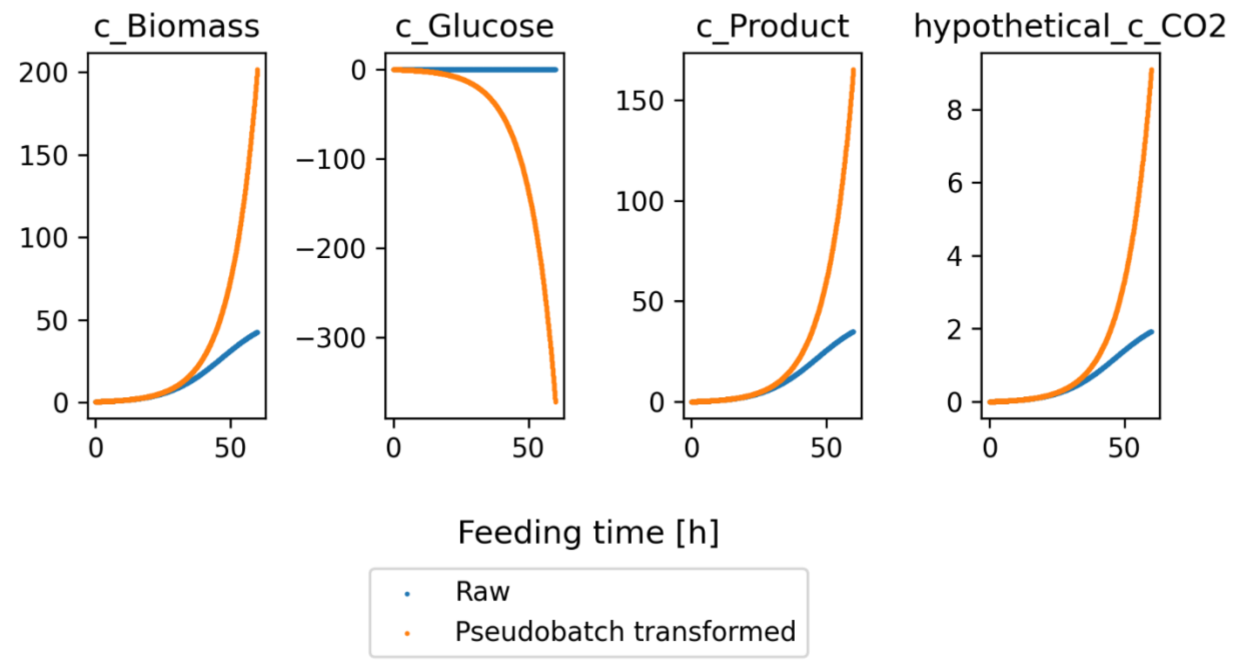

***Figure S2:*** *Raw simulated concentration data (blue) and pseudo batch transformed data (orange). For CO2 the plot shows the hypothetical concentration (see Hypothetical concentration) instead of the raw concentration data.*

***Table 1:*** *Parameters used to produce the simulated fed-batch process.*

| PARAMETER | VALUE | UNIT | PARAMETER DESCRIPTION |
|---|---|---|---|
| $K_s$ | 0.150 | g/L | The Ks value for the monod equation textbook estimates for *Saccharomyces cerevisiae* from [2] |
| $\mu_{max}$ | 0.300 | 1/h | The maximum specific growth rate textbook estimates for *Saccharomyces cerevisiae* from [2] |
| $Y_{xs}$ | 1.850 | gSubstrate / gBiomass | The yield of substrate on biomass textbook estimates for *Saccharomyces cerevisiae* from [2] |
| $Y_{xp}$ | 0.822 | gProduct / gBiomass | The yield of product on biomass textbook estimates for *Saccharomyces cerevisiae* from [2] |
| $Y_{xco2}$ | 0.045 | gCO2 / gBiomass | The yield of CO2 on biomass textbook estimates for *Saccharomyces cerevisiae* from [2] |
| $F_0$ | 0.063 | µL/h | The initial feed rate |
| $\mu_0$ | 0.100 | 1/h | The target specific growth rate used to calculate the feed rate |
| $c_s^f$ | 100.000 | g/L | The substrate concentration in the feed |
| **SAMPLE_VOLUME** | 170.000 | µL | The sample volume |
| **N_SAMPLES** | 8.000 | counts | The number of samples during the whole process |
| **DURATION** | 60.000 | h | The simulated duration of the fed-batch process |
| **X0** | 0.5 | gBiomass | Initial biomass concentration |
| **S0** | 0.075 | gSubstrate | Initial substrate concentration |
| **P0** | 0 | gProduct | Initial product concentration |
| **CO2_0** | 0 | gCO2 | Initial CO2 concentration |

#### S4 Estimation of overall rates and yields

We estimated the overall growth rate for the entire simulation period using a log-linear model in which the log-transformed biomass in either mass or pseudo batch concentration is a linear function of time.

$$\begin{aligned} \log\left(c_x^\star(t)\right) &= a + \hat{\mu} \cdot t \\ \log\left(m_x(t)\right) &= a + \hat{\mu} \cdot t \end{aligned}$$

where $c_x^\star(t)$ is the pseudo biomass concentration and $m_x(t)$ is the non-corrected mass at measurement at time $t$, and $\hat{\mu}$ is the fitted specific growth rate.

The overall yield coefficients, e.g. $Y_{sx}$, were estimated as the slope of a linear model between the measurements of the two species [3]. For comparison we also formulated the following two linear models for the uncorrected data:

$$\begin{aligned} cm_s(t) &= b + Y_{xs} \cdot m_x(t) \\ m_p(t) &= b + Y_{xp} \cdot m_x(t) \end{aligned}$$

where $cm_s(t)$ is the consumed mass of substrate at time $t$, calculated as $cm_s(t) = m_s(0) + \int_0^t v_{feed}\,(t) * c_f - m_s(t)$. For pseudo batch transformed data it is not required to calculate the consumed substrate. Thus, we used the following model to estimate overall yield coefficients from pseudo batch transformed data.

$$c_i^\star(t) = \mathrm{b} + Y_{xi} \cdot c_x^\star(t)$$

where $c_i^\star(t)$ is the pseudo batch concentration of species $i$ at time $t$. The parameter estimates of all the linear models were obtained as the ordinary least-squares solution using the Statsmodels Python package [4]. Uncertainties of the parameter estimates are represented using ± describe the standard error of the parameter estimate scaled by 1.96 to cover 95% confidence region.

We calculated the relative error of the parameter estimates through the following equation:

$$\text{Relative error } \% = \frac{\mathrm{P} - \hat{P}}{\mathrm{P}} * 100\%$$

Where $P$ the true simulated parameter, growth rate or yield coefficient, and $\hat{P}$ is the parameter estimate.

#### S5 Statistical model

We parameterized our statistical model in terms of the following unknown quantities:

| symbol | Description |
|---|---|
| $v_0$ | The volume of the container at the start of the experiment. |

| symbol | Description |
|---|---|
| $m$ | A matrix of species masses (one number per species per sample). |
| $\alpha$ | A vector of volumes, expressed as logit-scale fractions of the pre-sample volume (one number per sample). |
| $f_{nz}$ | A vector containing the feed volume for each pre-sample interval where feeding took place (one number per sample with non-zero pre-sample feed). |
| $cf_{nz}$ | The concentration of feed in the feed solution if any (one number). |
| $a_{pump}$ | The factor according to which measurements of feed volume are biased, on natural logarithmic scale (one number). |

Reactor volumes $v$ and sample volumes $s$ are derived from these unknowns as shown in this equation:

$$
\begin{aligned}
v_i &= \begin{cases} v_0 + f_1 & i = 1 \\ v_0 + \sum_{j=1}^{i} f_j - \sum_{k=1}^{i-1} s_k & i > 1 \end{cases} \\
s_i &= v_i \cdot logit^{-1}(\alpha_i)
\end{aligned}
$$

Species concentrations can then be calculated by dividing the masses $m$ elementwise by the volumes $v$.

This parameterization was chosen to allow for independent, semi-informative priors on the unknown quantities together with an intuitive measurement model, and to ensure that all model quantities obey known constraints, such as that volumes have to be non-negative.

Our model uses independent log-normal prior distributions for positive-constrained unknowns and independent normal prior distributions for unconstrained unknowns. To describe measurements, we used a log-normal generalized linear model with a natural logarithmic link function:

$$
\begin{aligned}
v_0 &\sim LN(\mu_{v_0}, \tau_{v_0}) \\
m &\sim LN(\mu_m, \tau_m) \\
\alpha &\sim N(\mu_\alpha, \tau_\alpha) \\
f_{nz} &\sim LN(\mu_{f_{nz}}, \tau_{f_{nz}}) \\
cf_{nz} &\sim LN(\mu_{cf_{nz}}, \tau_{cf_{nz}}) \\
a_{pump} &\sim N(\mu_{pump}, \tau_{pump}) \\
y_v &\sim LN(\ln(v), \sigma_v) \\
y_c &\sim LN(\ln(c), \sigma_c) \\
y_s &\sim LN(\ln(s), \sigma_s) \\
y_f &\sim LN(\ln(f + a_{pump}), \sigma_f) \\
y_{cf_{nz}} &\sim LN(\ln(cf_{nz}), \sigma_{cf_{nz}})
\end{aligned}
$$

The model specification is shown in mathematical notation in the equation below. Our model assumes that all measurement error terms, i.e. $\sigma_v$, $\sigma_c$, $\sigma_s$ and $\sigma_{cfeed}$ are known exactly. Each term $y_x$ refers to vectors of measurements of the corresponding quantity $x$, and the terms $N$ and $LN$ represent the normal and log-normal distributions respectively.

In our analysis we used semi-informative prior location and scale parameters based on quantiles, ruling out values that were implausible based on scientific knowledge. Users of our package can select these values according to the specific requirements of their analysis using our interface.

#### S5.1 Implementation

We specified our model using the Stan probabilistic programming language [5]. We fit this model to our simulated dataset using Stan's adaptive Hamiltonian Monte Carlo algorithm via the cmdstanpy interface [6] and used arviz [7] to store the resulting MCMC samples.

In order to confirm that our samples constituted an accurate enough summary of the target posterior distribution, we verified that there were no post-warmup divergent transitions, [8] that the improved $\hat{R}$ statistic was in the interval [0.99, 1.01] for all variables and that the estimated Monte Carlo standard error for transformed and untransformed concentration variables was orders of magnitude smaller than the precision with which we needed to estimate the uncertainty in these quantities as recommended in [9].

Code implementing our analysis, as well as instructions for reproduction, can be found at *https://github.com/biosustain/pseudobatch*

#### S5.2 Interface

In order to allow users to investigate the error propagation properties of their own experiments, we exposed the statistical model through the Python function `pseudobatch.run_error_propagation`. See the *documentation page* and the *pseudo batch with uncertainties tutorial* for information about how to use this function. In order to avoid the user having to install cmdstan, we used the template `cookiecutter-cmdstanpy-wrapper` [10] to bundle the model with our package.

#### S5.3 Why not do Monte Carlo sampling?

The reader might wonder why we constructed a full statistical model to investigate the error propagation properties of the pseudo batch transformation. An alternative approach would have been to construct only a prior model and then perform Monte Carlo sampling to empirically ascertain uncertainties.

Unfortunately, this approach was not feasible in our case because of the parameter correlations induced by the problem structure. Since reactor volumes, sample volumes, and concentrations depend on each other, any model must be parameterised in terms of auxiliary parameters as described above in the section on *Implementation*. However, there is typically information available about reactor volumes, sample volumes and measurements. In a full statistical model this information can be account for with a measurement model. In a priors-only approach as required for Monte Carlo sampling, this information must be represented by imposing a corresponding covariance structure on the auxiliary parameter priors. We judged that this would be prohibitively difficult, so we created the full statistical model instead.

#### S5.4 Case study

To test our model, we used the plausible simulated fed-batch dataset described above in the *Simulated data* section and carried out the steps described in the *Implementation* section, with a measurement error on all the concentrations ($\sigma_c$) set to 0.05. We plotted the 1% to 99% marginal posterior interval for each species to compare the pseudo batch transformed and untransformed time courses:

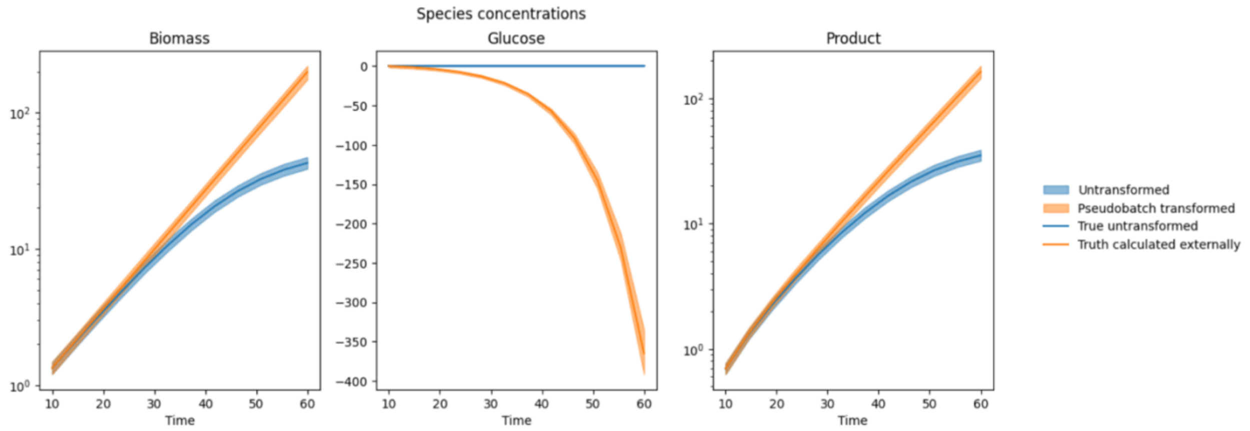

***Figure S3:*** *Concentration time times of the biomass, glucose and product. The blue line shows the true simulated time courses, and the orange line is the pseudo batch transformed true time course. The shaded areas span the 1% to 99% credible interval of the concentration time series as calculated from the Bayesian model using* $\sigma_c = 0.05$.

Next, we varied the input $\sigma_c$ in 30 increments between 0.01 and 1 and compared the marginal posterior coefficients of variation of pseudo batch transformed biomass and product concentration with that of the corresponding untransformed concentrations (glucose was excluded here because it was simulated as present in the feed). The results are shown below:

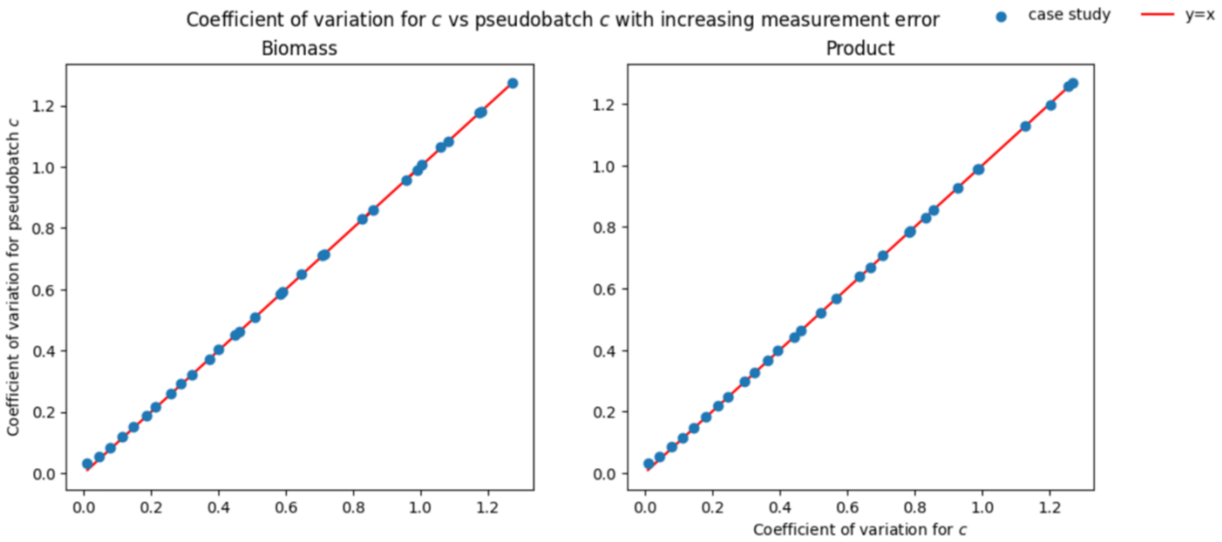

***Figure S4:*** *Coefficient of variation in the pseudo batch transformed concentration values compared to the input measurement error (as coefficient of variation). The red line denotes the trajectory where the two coefficient of variation values are exactly equal, i.e. when the transformation does not change the measurement error.*

These results show that for the full range of measurement error assumptions, the pseudo batch transformed variable has approximately the same posterior coefficient of variation as the untransformed variable.

Finally, we estimated the growth rate and product and glucose yield coefficients for each posterior sample using the method described in section *Estimation of overall rates and yields*. The results are shown below alongside the true values used in the simulation in figure S5.

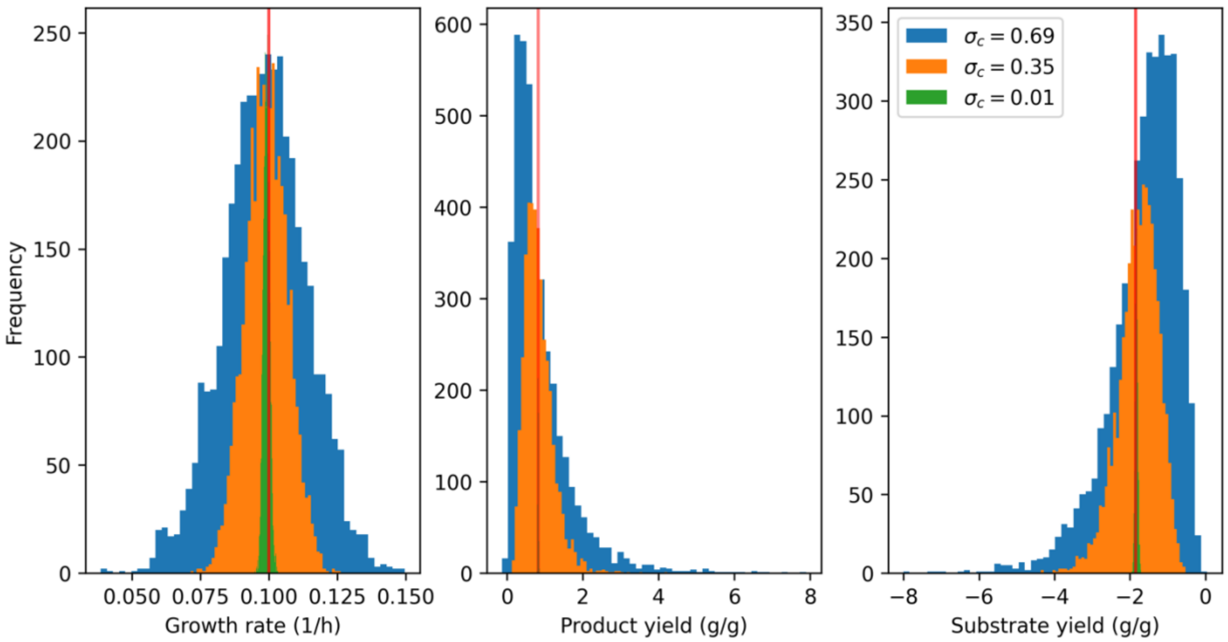

***Figure S5:*** *Distributions of parameter estimates from posterior samples using increasing measurement error ($\sigma_c$). The true value of the parameter is shown as a red line.*

These histograms are not exactly centered on the true values (red line), thus there is a small consistent bias compare the true values. This bias originated from prior model. As the measurement uncertainty increases the posterior distribution is pulled towards the prior distribution. Table S3, show how the 50% quantile of the posterior distributions moved towards the 50% quantile of the prior predictive distribution as $\sigma_c$ increases.

***Table S3:*** *50% quantiles of the parameter estimates based on posterior samples from different measurement errors ($\sigma_c$'s) and estimated from the prior predictive distribution.*

| DISTRIBUTION | GROWTH RATE | PRODUCT YIELD | SUBSTRATE YIELD |
|---|---|---|---|
| POSTERIOR WITH $\sigma_c$ = 0.69 | 0.098608 | 0.644731 | -1.434936 |
| POSTERIOR WITH $\sigma_c$ = 0.35 | 0.099104 | 0.761890 | -1.705882 |
| POSTERIOR WITH $\sigma_c$ = 0.01 | 0.099141 | 0.821709 | -1.825006 |
| PRIOR PREDICTIVE | 0.026289 | -0.012052 | -0.019035 |
| TRUE VALUE | 0.10 | 0.82 | -1.85 |
